# To call or not to call: Persistence of flexible alternative reproductive tactics in a tree cricket

**DOI:** 10.1101/2023.04.30.538844

**Authors:** Mohammed Aamir Sadiq, Viraj R. Torsekar, Rohini Balakrishnan

## Abstract

Alternative reproductive tactics (ARTs) are discrete polymorphisms that help maximise reproductive success. Although flexible ARTs are ubiquitous, theoretical predictions for why flexible ARTs persist over evolutionary time have rarely been empirically tested. We hypothesised that flexible ARTs will persist if they have equal fitness benefits under a range of ecological contexts, or, there are trade-offs between ARTs in different ecological contexts and individuals display the most optimal phenotype in a context-dependent manner. Specifically, we investigated predation risk effects on the expression and fitness consequences of two flexible ARTs: acoustic signalling and being silent, expressed by tree cricket *Oecanthus henryi* males. In large outdoor enclosures, we exposed natural populations of *O. henryi* to three different abundances of their predator, the green lynx spider *Peucetia viridans*. Behavioural observations across successive nights revealed that higher predation risk did not alter the expression levels of the male ARTs, despite crickets experiencing differential risk and survival across treatments. Male crickets demonstrated an equal likelihood of calling or remaining silent on a night. ARTs resulted in similar mating success across differential predation risk, supporting the hypothesis that equal fitness benefits of flexible ARTs under a range of ecological contexts explain their persistence.

## Introduction

The origin and persistence of alternative reproductive tactics (ARTs) has been a subject of broad interest to evolutionary biologists. ARTs are discrete phenotypes that help maximise reproductive success in the context of sexual selection (Brockmann & Taborsky, 2008). They are extensively found in sexually reproducing organisms (Brockmann et al., 1979; Brockmann & Taborsky, 2008; Howard, 1978; Lank et al., 1995) and provide unique opportunities to study the evolution and maintenance of phenotypic variation, a key focus of evolutionary biology. ARTs can be genetically polymorphic or plastic (Gross, 1996). While genetically polymorphic ARTs are fixed for an individual’s lifetime, plastic ARTs can be either fixed or flexible, that is, individuals can switch between ARTs during their lifetime. For example, based on their nutritional states, males of the scarab dung beetle (*Onthophagus acuminatus*) express different morphs which remain fixed for the remainder of their lifetime (Emlen, 1997). In contrast, males of the desert grasshopper (*Ligurotettix coquilletti*) can reversibly switch between calling and satellite tactics across nights (Greenfield & Shelly, 1985).

Theory suggests that while genetically polymorphic ARTs can persist in a population only when their average fitnesses are equal (Shuster, 2010), plastic ARTs can persist whether they have equal fitnesses or not (Tomkins & Hazel, 2007). While evidence suggests that plastic ARTs may not have equal average fitness (Brockmann et al., 1994; Ferrari et al., 2019; Hunt & Simmons, 2001; Wilgers et al., 2009), the conclusion is premature because field studies that estimate average fitnesses of ARTs tend to exclude unmated males, which inflates the mean and underestimates the variance of average fitness (Shuster, 2010). Furthermore, most studies evaluating fitness of ARTs have focussed on fixed ARTs, with little attention paid to flexible ARTs (but see Brockmann & Penn, 1992). Understanding the fitness of flexible ARTs will further our understanding of the selection regimes under which fixed and flexible plastic ARTs evolve.

Flexible ARTs can theoretically persist if they incur similar costs and benefits across a range of ecological contexts, leading to equal fitness (Shuster, 2010). Individuals may, however, frequently switch between ARTs at short time scales. An extreme example of flexible ARTs can be found in males of a marine bioluminescent ostracod (*Photeros annecohenae)* which can switch among three different photo-signalling patterns within 12 seconds (Rivers & Morin, 2009). Such dynamic switching between ARTs may be a response to maximising fitness in a frequently changing environment. For example, in the ecological context of increased predation risk, a conspicuous sexual signalling tactic could attract predators and experience elevated costs (Johnson & Candolin, 2017). An inconspicuous ART, however, could substantially reduce the probability of being preyed upon (Clifton & Robertson, 1993). Although individuals expressing such inconspicuous ARTs will encounter limited number of mates, the fitness trade-off in this risky context would favour selection of the phenotype associated with it (Rotenberry & Zuk, 2016; Walker & Cade, 2003). Thus individuals are expected to express the most optimal phenotype, given the trade-offs in fitness between the ARTs in the appropriate ecological context (Lively, 1986). The average fitness of the optimal phenotypes may be unequal but they will be maintained provided there is sufficient variation in ecological contexts (Hazel et al., 1990).

Measurement of fitness consequences of flexible ARTs requires collecting longitudinal information on the number of times individuals in a population express each of the flexible ARTs, as well as the benefits and costs associated with each expression to demonstrate average payoffs of the ARTs for each individual. These observations have rarely been made in natural populations (but see Brockmann et al., 1979). Brockmann et al. (1979) performed longitudinal observations on the flexible nesting behaviour of female digger wasps (*Sphex ichneumoneus*) and found that the pay-off obtained by females from adopting nest parasitism was comparable to that obtained from nest-building. They, however, calculated the payoff of the ARTs at the population level, whereas individual payoffs are a more pertinent measure to empirically examine the theoretical requirement of fitness equality or inequality of ARTs.

A commonly studied ART, sexual signalling and mate searching behaviour, generally involves signalling and searching for mates, and is prone to increased predation risk (Sih, 1994). Predation risk is an important ecological context that can affect i) level of expression, and ii) consequent fitness of ARTs (Rotenberry & Zuk, 2016; Walker & Cade, 2003). Under risky conditions, individuals may exhibit high repeatability in tactics that increase the likelihood of survival (Toscano et al., 2014) and may switch to less conspicuous ARTs (Sih, 1994). For example, male guppies shift from showing conspicuous sigmoid displays to adopting a sneaker tactic when presented with a model predator (Godin, 1995). Likewise, *Photinus* fireflies alter their display patterns during courtship in the presence of visually orienting predators (Woods et al., 2007). Predation risk can also influence the mating success of ARTs by altering the reproductive behaviour of receivers of signals (Johnson & Basolo, 2003). If signalling males attract predators, females are expected to avoid such males due to the predation risk associated with them and may also be less choosy (Sih, 1994). While some studies have focused on change in expression of ARTs when exposed to predators (reviewed in Bernal & Page, 2022), the relationship between expression of ARTs and their corresponding fitness effects under varying predation risk has received little attention.

In this study, we aim to link effects of predation risk on expression patterns of flexible ARTs with their resulting fitness consequences under semi-natural conditions. We used the tree cricket (*Oecanthus henryi*) and its major predator, the green lynx spider (*Peucetia viridans)* as our model system (Torsekar et al., 2019). *Oecanthus henryi* exhibits a mating system typical of crickets, where males produce species-specific calls and females use the calls to localise conspecific males (Walker, 1957). Alternatively, males can switch to a silent tactic on some nights, observed commonly in many other taxa (Loher & Orsak, 1985; Woolbright, 1985). Silent cricket males may eavesdrop on calling males and behave as satellites to intercept females (Cade, 1981; Zuk et al., 2006). To examine the flexibility of ART expression in *O. henryi*, we quantified repeatability of ART expression. Flexible switching between calling and silent tactics raises the possibility that males may also use the silent tactic to mate.

We considered calling and silent as two flexible ARTs that can be expressed by males to investigate how environments with differential predator abundance affect i) male calling propensity (probability of calling on a given night) and ii) mating success of the flexible ARTs. We predicted that males would reduce their calling propensity under higher predator abundance due to perceived risk. Furthermore, given that both ARTs coexist in natural populations, we predicted that the mating success of these ARTs will follow one of two predictions in response to predator abundance. First, the mating success of both ARTs may be similar across different levels of predation risk, thereby supporting their co-existence (Fig. 1A). This scenario can arise when females do not exhibit mate choice based on male signals across multiple ecological contexts. Although female crickets generally prefer louder calls (Mhatre & Balakrishnan, 2007; Deb, et al., 2020), females in their natural habitat do not perform mate sampling and mate with the first male they encounter, which may not necessarily be the louder one (Nandi et al., 2019). Further, females will mate with the first male they are presented with, in the absence of song, in laboratory experiments (Modak et al., 2021). Alternatively, fitness trade-offs could be associated with ARTs at different predator abundances. Lower predator abundance, by lowering the risk of mortality, will allow females to exercise mate choice and choose calling males over silent ones, while at higher predator abundances, females may lower their risk of mortality by avoiding calling males they perceive to be at higher risk and mating with the less conspicuous silent males. If that is the case, we hypothesised that the calling tactic would have higher mating success at lower predator abundances compared to the silent tactic and vice versa at higher predator abundances (Fig. 1B).

**Figure 1.**
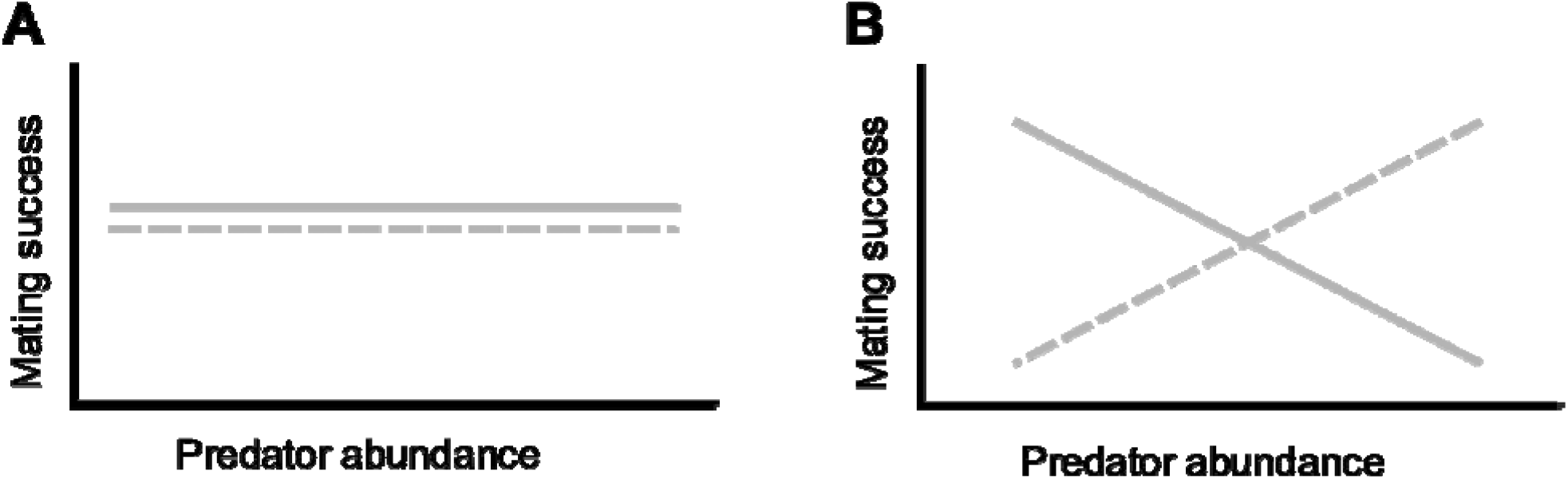
Graphical illustration of predicted effect of predator abundance on mating success of alternative reproductive tactics (ART) expressed by male crickets. Solid line indicates mating success per expression of the calling tactic and dashed line indicates mating success per expression of silent tactic. Given that both ARTs coexist in natural populations, two explanatory scenarios are possible: A) Mating success of both ARTS per expression are similar at all levels of predator abundance possibly due to the absence of mate choice by females, or B) Mating success of both ARTS show trade-offs with respect to predator abundance. Here, calling tactic is more beneficial at low predator abundances possibly due to females preferring calling males. Silent tactic is beneficial at higher predator abundance possibly due to female avoidance of calling males who could attract predators.

## Methods

### Field experiment

We conducted our study in outdoor enclosures (6m × 6m × 3m) near Ullodu (13°38′27.2″N 77°42′01.1″E) village, in fields of the Chikkaballapur district of Karnataka state, India, from February 2016 to May 2017. The enclosures were constructed using a stainless-steel mesh supported by wooden pole scaffoldings around naturally occurring, homogeneously distributed *H. suaveolens* bushes (Torsekar & Balakrishnan, 2020). The vegetation inside the enclosures was representative of the surrounding field site where the enclosures were located. We conducted three different predation treatments that differed in the predator-prey ratios introduced in the enclosures: ‘No predation’, ‘Low predation’ and ‘High predation’. Fifteen *O. henryi* males and fifteen females were released in an enclosure for each of the three treatments. In the ‘No predation’ treatment, no spiders were introduced into the cage. The ‘Low predation’ and ‘High predation’ treatments involved releasing 15 and 120 spiders into the cage, respectively. Thus, the predator: prey ratio for the three treatments were as follows: No predation (0:30), Low predation (15:30) and High predation (120:30). The adult spiders and crickets were wild-caught two days prior to the start of each trial from the field site. We marked all crickets individually with distinct colour codes using non-toxic paint markers (Edding 780) to facilitate individual identification during the trial. Crickets were released on randomly chosen bushes in the enclosure 24 hours prior to the trial to allow the crickets to acclimatise. One hour before the trial, we released the spiders on randomly chosen bushes.

*Oecanthus henryi* exhibits peak activity between 7.00 pm and 10.00 pm (Deb & Balakrishnan, 2014). Consequently, we recorded the behaviour and location of crickets from 7.00 pm to 9.30 pm. We used the instantaneous scan sampling method to record the location and behaviour of individual crickets. The location of tree crickets was recorded thrice every night at one-hour intervals, i.e., at 7 pm, 8 pm and 9 pm. We also recorded calling and mating occurrences for all male crickets every 10 minutes. Thus, we made 16 behavioural scans every night that provide us with data pertaining to calling and mating activity of individual male crickets. We did not replace crickets that died during the trial. However, as the number of crickets in the cage reduced during the trial, we removed spiders accordingly to maintain a constant predator: prey ratio. For each trial, we sampled an enclosure for 10-12 continuous nights. The high and low predation treatments were replicated thrice, and the no predation treatment was replicated twice. Thus, we conducted eight trials in total.

To ensure that the experimental manipulation was effective, we measured the predation risk experienced by all crickets across all nights they survived for each treatment. We used spatial proximity between crickets and spiders as a proxy for predation risk by quantifying the probability of them co-occurring on the same bush. For each individual cricket, we calculated predation risk as the proportion of sampled nights it was observed to co-occur with a spider on the same bush. Differences in predation risk experienced by individual crickets across the three predation treatments were estimated using permutation tests (details in S1). Survivorship differences of crickets as a consequence of exposure to the three predation treatments was estimated using the Kaplan-Meier method and confidence intervals were plotted to visually inspect differences between the survival curves (Kaplan, E. L., and Meier, 1958). The survivorship curves were constructed using the ‘ggsurvfit::survfit2’ R package (Sjoberg and Baillie, 2022).

### Classification of males into caller and silent ARTs

We used call effort (proportion of behavioural scans out of a total of 16 on a night in which a male was observed to be calling) to arrive at our definitions of ARTs. The histogram of call efforts of all males pooled across all nights obtained from replicates of all treatments indicated a clear mode at 0 (supplementary figure S1), suggesting that being silent on a night was a frequent tactic exhibited by males. Therefore, we defined a silent ART as a male that never called for the sampled duration of a night. Alternately, we defined a calling ART as a male that had a calling effort greater than 0 for a night.

### Flexibility of ARTs

Individual males of *O. henryi* can potentially switch between calling and silent ARTs across nights, showing flexibility in the expression of ARTs. We quantified the flexibility in expression of calling within an individual in terms of repeatability. Repeatability is a common measure used to quantify the constancy of a phenotype (Nakagawa & Schielzeth, 2010). Since the relevant ARTs are of two types, calling and silent, we scored the expression of ARTs by a male on an individual night as a binary variable (0 for silent and 1 for calling male). Consequently, we estimated the repeatability of the expression of these binary ARTs across nights using the rptBinary function in the R package ‘rptR’ (Nakagawa & Schielzeth, 2010). Since the rptBinary functions uses a default logit-link, the model estimates are fitted on the latent scale. We calculated the latent-scale repeatability under each predation treatment separately after lumping data across replicates for the relevant treatment. Individual male ID was used as the grouping variable. We used bootstrap resampling (N = 1000) to obtain 95% confidence intervals for each repeatability estimate.

### Calling Propensity

To quantify the decision of males to exhibit one ART over the other, we measured the probability of expression of the ARTs in terms of calling propensity. Calling propensity is the tendency of a male to call on a night. We calculated the propensity of a male to call on a night as the ratio of the total number of nights a male called to the number of nights the male was observed. Thus, calling propensity is essentially the probability of expression of calling on a night for a male.

We then tested whether the probabilities of expression of the two ARTS are similar under each predation treatment. For each predation treatment, we compared calling propensity to the assumed null median of 0.5 using a Wilcoxon signed-rank test. A deviation from 0.5 would indicate that a male is more likely to either call or remain silent on a night. Further, we performed a Kruskal-Wallis test (Hollander et al., 2013) to compare call propensities across predation treatments to check if predator abundance affected the choice of ARTs exhibited by the males.

### Mating Success

We used mating success as a proxy to evaluate the fitness payoffs of the ARTs. Mating success for each ART was measured as the ratio of the total number of matings obtained by a male when performing a tactic to the number of nights the male performed that particular tactic. Thus, mating success is essentially the payoffs a male receives for each expression of an ART (calling or silent on a given night). To analyse the effect of predation on the mating success of ARTs, we performed a Poisson Generalised Linear Model (GLM) using loglink (McCullagh & Nelder, 1989), with predation treatments, type of ARTs and their interaction as fixed factors. Since mating success in our analysis could be a fraction, we considered the number of matings by an individual male while expressing a type of ART as the response variable and used the number of nights for which the male expressed the ART as the offset term. All analyses were performed on R software version 3.4.3 (R Core Team, 2022).

## Results

### Predation risk and Survivorship

Crickets experienced different predation risk in each treatment since the probabilities with which crickets co-occurred with spiders were different across treatments (p < 0.001; Fig. 2). We observed a significant difference in predation risk experienced by both males and females across any two given predation treatments, when sexes were analysed separately (p < 0.001 for both males and females). Lastly, there was no significant difference between the predation risk experienced by males and females in any given predation treatment (No predation: p = 1, low predation: p = 0.6378, and high predation: p = 0.602). Survivorship was lower in high predation treatments in comparison with no and low predation treatments, for both males (Fig. 3A) and females (Fig. 3B).

**Figure 2.**
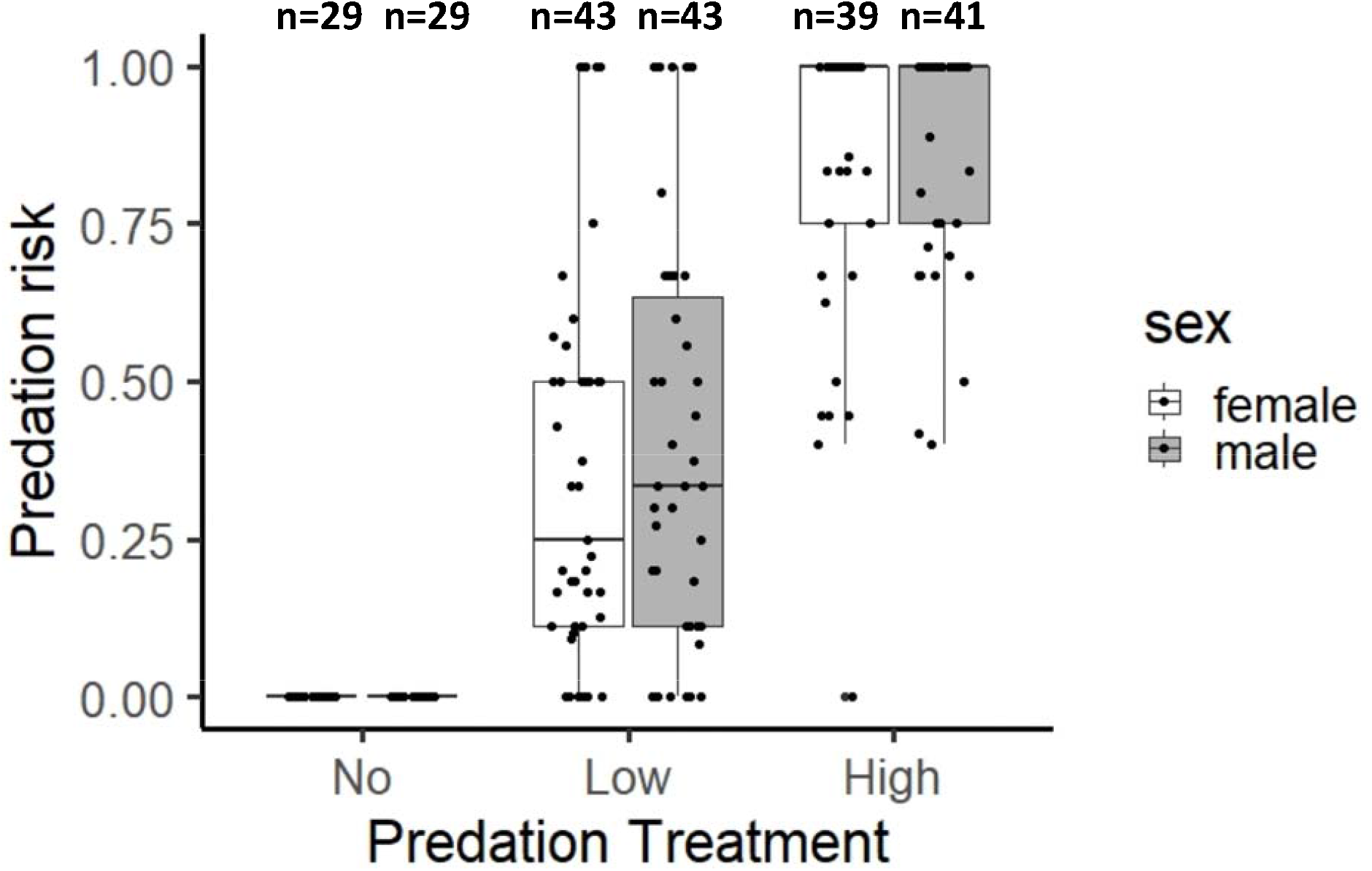
Predation risk experienced by male and female crickets across the three predation treatments. Predation risk, measured as probability of co-occurrence of crickets and spiders on a bush, is different across the treatments, but similar for both sexes within a treatment.

**Figure 3.**
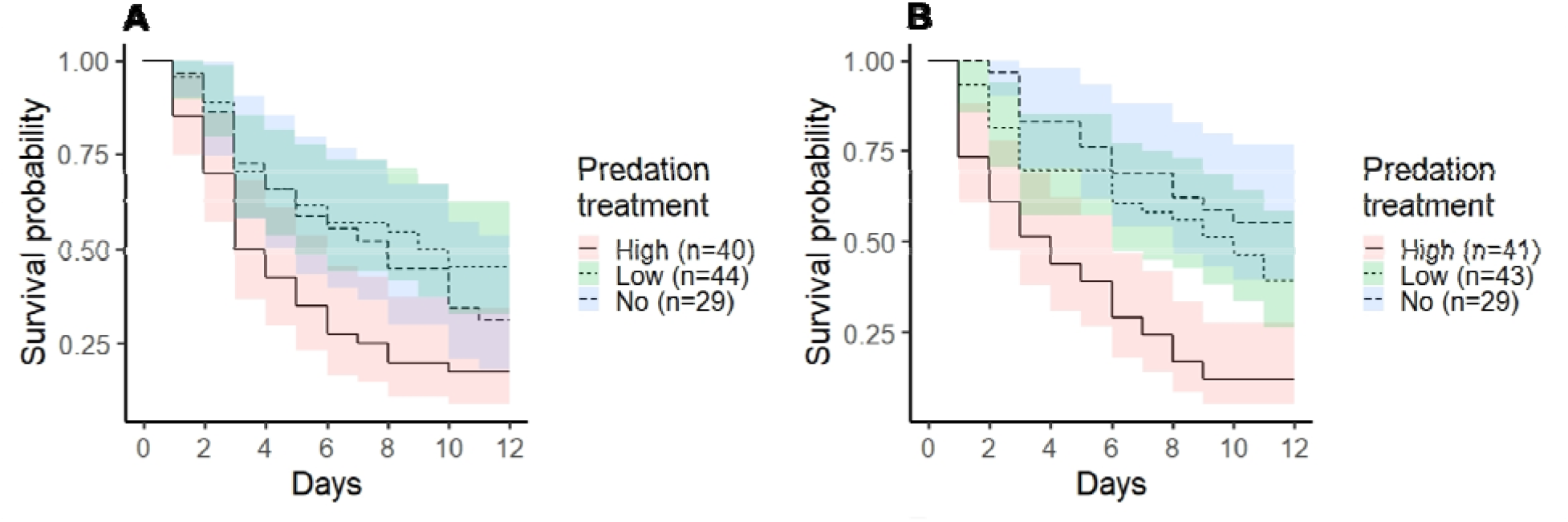
Survivorship curves of A) males and, B) females under the three predation treatments show that survivorship was lower in high predation treatments in comparison with no and low predation treatments, for both males and females. Shaded regions indicate 95% confidence intervals.

### Flexibility of ARTs

The latent-scale repeatability of across-night expression of ARTs in each of the predation treatments was, no predation: 0.336 (95% CI = 0.109 to 0.553, N = 29), low predation: 0.307 (95% CI = 0.145 to 0.480, N = 44), and high predation : 0.362 (95% CI = 0.147 to 0.605, N = 40). The high overlap of the 95 % CI between the predation treatments suggests that the across-night expression of ARTs was similarly flexible across the predation treatments (Fig. 4A).

**Figure 4.**
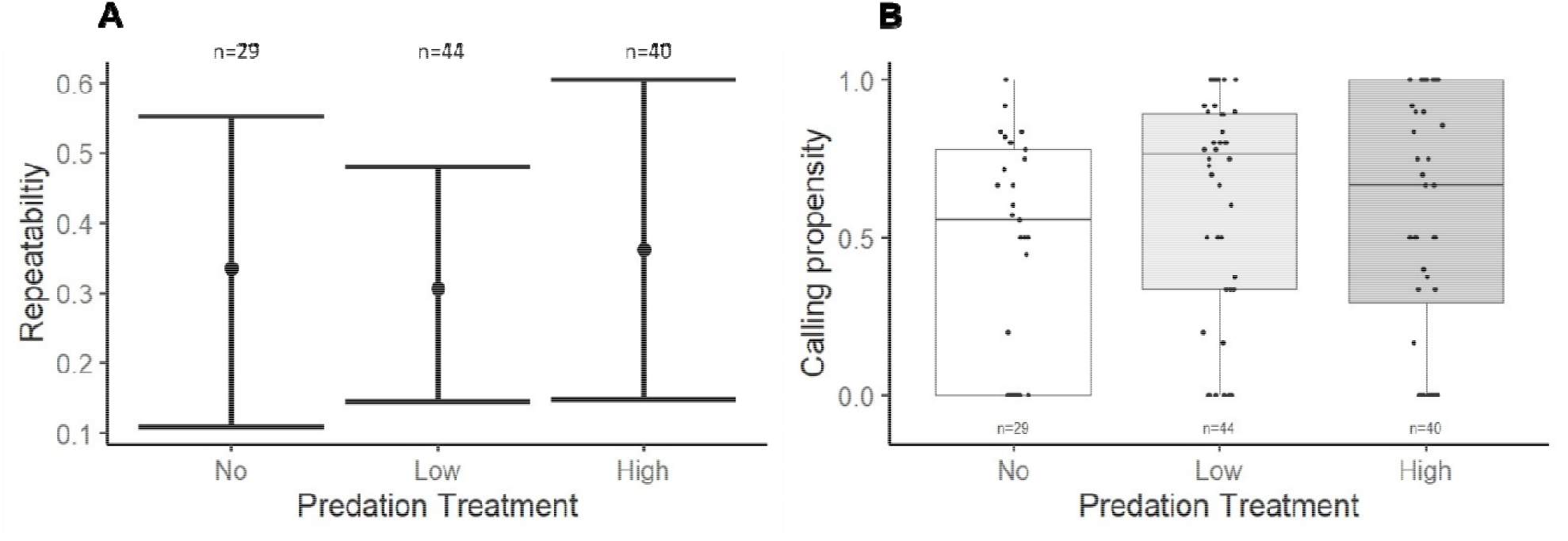
Expression of alternative reproductive tactics (ARTs). A) Across-night repeatability of expression of ARTs (silent and calling) across predation treatments. Error bars indicate 95% confidence intervals obtained after bootstrap resampling 1000 times. Sample size n indicates the number of individual males from which the repeatability estimate was calculated for each treatment. B) Calling propensity of individual males across the three predation treatments. Data were pooled across replicates for each treatment.

### Calling Propensity

Male calling propensity was found to be similar across the three predation treatments (Kruskal Wallis H test, p = 0.15), suggesting that predation did not affect the level of expression of the silent or calling tactics on a night (Fig. 4B). Furthermore, when we applied the Wilcoxon test to each predation treatment, call propensity was not significantly different from 0.5 in all three predation treatments (No predation, p = 0.45; Low predation, p = 0.07; High predation, p = 0.31). Males were thus equally likely to call or remain silent on a night irrespective of predator abundance.

### Mating Success

The results from the generalised linear model (Table 1, Fig. 5) indicate that the mating success of the silent tactic was not significantly different from the calling tactic (95% CI = -3.123 to -0.011, p-value = 0.09). Furthermore, mating success of either tactic in the high predation treatment was comparable to that in low and no predation treatments (95% CI of coefficient of low predation = -0.411 to 0.839, p = 0.54; 95% CI of coefficient of no predation = -0.849 to 0.687, p = 0.85), implying that mating success of both ARTs were similar across predation treatments. Lastly, there was no significant interaction of silent tactic with the low (interaction term: p = 0.12) and no predation (interaction term: p = 0.08) treatments (Table 1). The similarity in mating success between the ARTs, the similarity in mating success of either ART across predation treatments and the lack of significant interaction between ARTs and predation treatments together imply that the similarity in the per-capita mating success of the two ARTs was consistent across predation treatments.

**Table 1.**
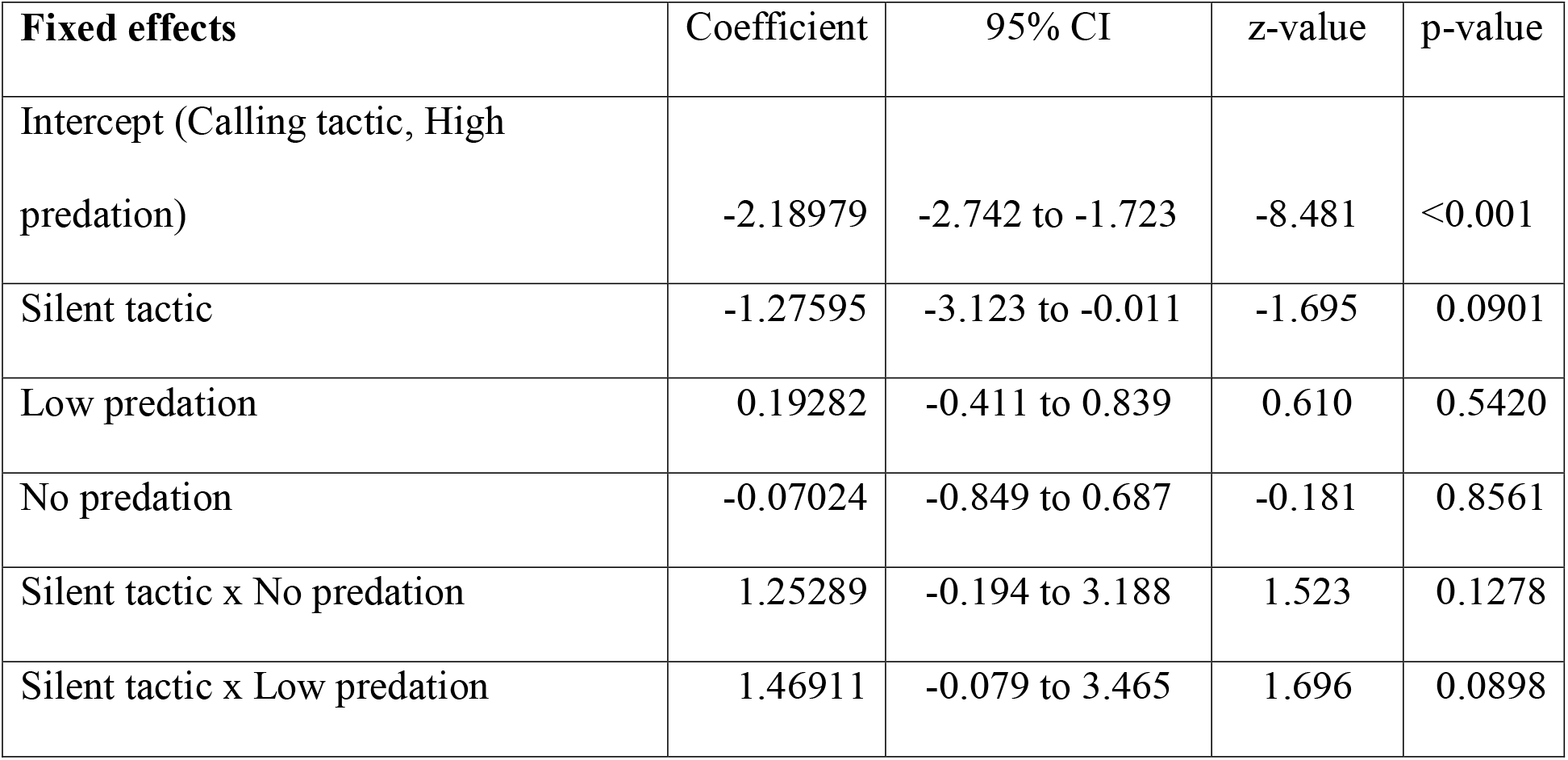
Coefficients of the predictors from the Generalised Linear Model used to analyse per-capita mating success of ARTs across predation treatments. The intercept represents coefficients of mating success associated with calling ART in the high predation treatment. Significance level was α=0.05.

**Figure 5.**
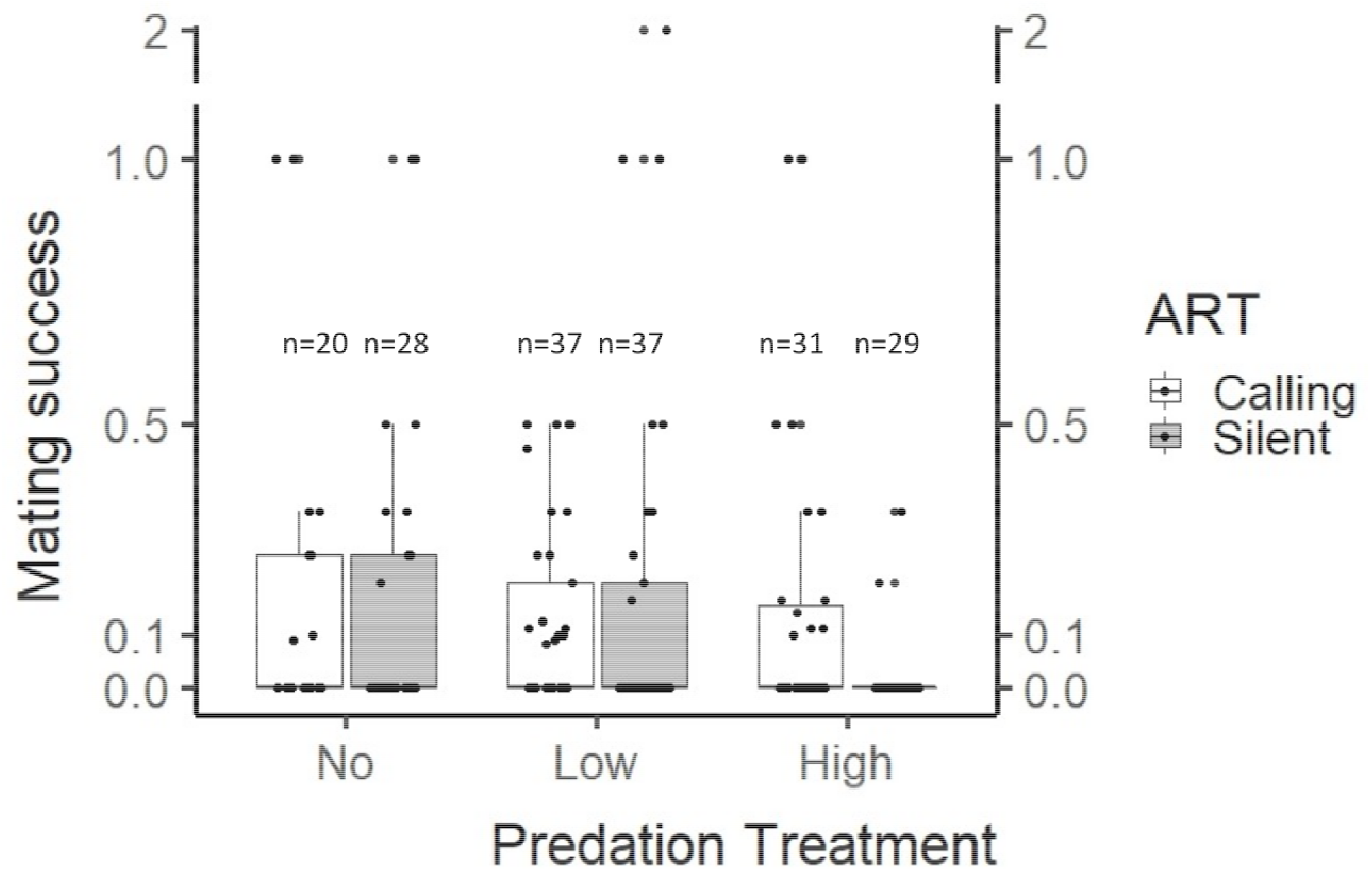
Mating success per expression of an alternative reproductive tactic (ART) for individual males. Each black dot represents the ratio of the total number of matings obtained by a male while expressing an ART to the number of nights the male expressed that ART.

## Discussion

The study of the persistence of flexible ARTs and the modulation of their fitness effects by the environment has hitherto received little empirical treatment. We examined the expression and fitness benefits of flexibly expressed ARTs, calling and silent, in wild-caught males of a tree cricket species in semi-natural conditions under different predation treatments. We found that males were equally likely to call or remain silent on a night irrespective of predator abundance. To estimate fitness consequences of the flexible ARTs, we examined mating success associated with each ART and found that both calling and silent ART elicit similar numbers of matings per night.

### What factors drive expression of calling behaviour?

The propensity to express ARTs did not differ in male crickets despite experiencing different levels of predation risk (Fig. 2) and differential survival (Fig. 3). Importantly, *O. henryi* crickets perceive the presence of a spider on the same bush as risky since they alter their within-bush movement patterns in the presence of the spider (Torsekar & Thaker, 2020). Therefore, similar calling propensity across predation treatments suggests that calling may not manifest as an additional survival risk from predation by the green lynx spider. This is in agreement with the fact that tree crickets experience similar probability of mortality from spider predation while calling and being silent (Torsekar et al., 2019). Under highly risky conditions, however, males reduce their calling effort in the presence of predatory spiders in their neighbourhood (Torsekar & Balakrishnan, 2020). These observations can be reconciled by the fact that initiation of calling on any night is nutrition-dependent (Modak, 2021) whereas how long crickets call may be modulated by more immediate factors such as perceived predation risk (Torsekar & Balakrishnan, 2020). Laboratory experiments with the same tree cricket species have demonstrated that males alter their within-night calling effort and nightly calling propensity in response to nutrient manipulations (Modak, 2021). In fact, males kept on a natural diet of *H. suaveolens* leaves showed a calling propensity of ∼0.5, 10-15 days post eclosion (Modak, 2021), similar to our results under varying predation risk, strengthening our conclusion that calling propensity is more likely driven by nutritional resources than by predation risk.

### How do flexible ARTs co-exist?

Persistence of flexible ARTs across evolutionary time is typically attributed to equal relative fitness of each ART across a range of ecological conditions. We evaluated the mating success of ARTs under a gradient of predator abundances and found that the mating success of ARTs was equal. The absence of any significant difference in the payoffs of the two tactics despite a steep gradient of predator abundance implies that both ARTs are equally effective at acquiring mates, regardless of the predation risk. These results are in agreement with our prediction that equality of fitness in flexible ARTs across a range of risky conditions will ensure their persistence and in contrast to the prediction that the two ARTs will show trade-offs as a function of predator abundance. Past work in tree crickets has shown that even under varying risk conditions, females exhibit similar likelihood of movement across bushes (Torsekar & Balakrishnan, 2020), indicating that predator presence has little impact on female mate searching behaviour. Such consistency in female searching behaviour may result in similar encounter rates between males and females regardless of predation risk, leading to similar male mating success across predation treatments.

However, this alone may not explain similar mating success between ARTs since laboratory experiments have shown that non-mated females are highly attracted towards conspecific male calls (Modak et al., 2021). Only about 30% of wild-caught females, however, exhibited phonotaxis (Torsekar, 2020). Thus, the preponderance of non-phonotactic females in natural populations may partially reduce the mating success of the calling ART. Despite their lack of motivation to approach calling males, however, mated females willingly re-mate if a male is nearby (Modak et al., 2021), which is likely to occur if males and females co-occur on a bush. Therefore, encounters between mated females and calling or silent males may be equally probable and lead to similar mating success. Moreover, movement patterns of calling and silent males within a bush are similar regardless of presence of a predator on the bush (Torsekar & Thaker, 2020), lending support to the argument of similar male-female encounter rates for both calling and silent males. A closer analysis of movement and space use of individuals in their natural habitat will help highlight the mechanistic underpinnings of male-female encounter rates and their dependency on the male ARTs.

### Are silent males adopting satellite strategies?

In our study, silent males could have inflated their mating success by behaving as satellites of calling males. However, only 17% of matings involving silent males across all predation treatments occurred when a calling male was in the vicinity, that is, present on the same bush as the silent male(s). This suggests that satellite behaviour may not be a more beneficial strategy for silent males to adopt to obtain mates. While a study on male bullfrogs (*Rana catesbeiana*) has shown that the cost of being a satellite may be less than that experienced in territorial competition, and a model suggests gains in mating benefits due to the satellite strategy (Howard, 1978; Waltz, 1982), there is a dearth of empirical evidence to show that the satellite strategy is indeed more beneficial for silent males. Tractable model systems and manipulative experiments may provide insights into the potential benefits of satellite behaviour.

### Why should males call at all?

If calling and silent ARTs are equally effective at obtaining mates, individuals should shift to being silent to avoid the energetic costs of calling (Hoback & Wagner, 1997). However, calling males may offset the energetic costs by availing a first-male fertilisation advantage. Female crickets may mate multiply and can store sperm from different males (Ono et al., 1995). Male reproductive success may be positively correlated to the proportion of sperm it contributes to the spermatheca (Simmons, 1987). For example, non-mated tree cricket females are highly phonotactic and show higher spermatophore attachment duration (SPAD) with the first male they mate with (Modak et al., 2021). The high phonotaxis propensity of virgin females along with higher SPAD associated with first matings may allow calling males to attract and mate longer with virgin females, increasing their sperm transfer (Brown, 1997) and consequently improving reproductive success, thereby driving the persistence of calling as an ART in the population. Paternity analysis using micro-satellite markers may elucidate the relationship between first male mating, sperm transfer and reproductive success, helping us gain insights into whether first male mating indeed has an advantage.

### Conclusion

To summarise, our mesocosm experiments demonstrate that flexible ARTs that are expressed in equal measure by individual males in different ecological contexts also receive similar mating benefits. More such tractable mesocosm experiments on other systems expressing flexible ARTs across relevant ecological contexts, for example territory quality and foraging opportunities, will help test the generality of the equal fitness hypothesis for persistence of flexible ARTs.

## Author contributions

MAS, VRT and RB conceived the study and designed the experiments. VRT collected data and MAS analysed the data. MAS wrote the first draft of the manuscript and all authors contributed to interpretation of results and the writing of the manuscript.

## Acknowledgements

We thank Kada Reddy and Manjunatha Reddy for their help in constructing the enclosure and Prakash, Chandra, Nanjundaiah and Manjunatha for their assistance in fieldwork. The research was funded by DBT-IISc Partnership Programme (phase 1 and 2) of the Department of BioTechnology (DBT), Government of India and some of the equipment used in the study were funded by DST-FIST (Department of Science & Technology Fund for Improvement of S & T Infrastructure, Govt. of India). The research fellowship of MAS was funded by the Prime Minister’s Research Fellowship, Government of India. The authors have no conflict of interest to declare.

