## Supplemental information for "To call or not to call: Persistence of flexible alternative reproductive tactics in a tree cricket"

**Supplementary Material**

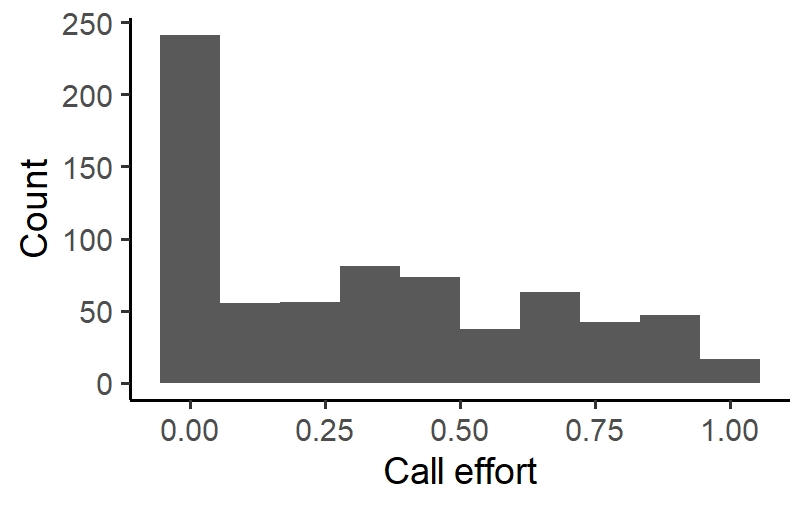

**Figure S1.** Histograms of call efforts of males pooled across all treatments and across all nights (n=711). A clear mode appears at 0 (Frequency=241) suggesting that being silent is a frequent ART exhibited by males. Males with calling effort greater than 0 were considered as calling males.

**S1. Permutation test**

We performed permutation tests to compare the predation risks of males and females across different predation treatments. In a permutation test, the absolute difference between the observed means (of the variable of interest) of the two groups of interest is calculated. Next, the data of the two groups are pooled and randomly sub-setted into two groups. The sizes of the randomised groups is equal to those of the original two groups of data. The absolute value of the difference of the means of the two groups of randomised data is calculated. Many iterations of randomising and sub-setting of the original data are done to obtain a null distribution of the absolute values of the differences of the means of the two randomised groups. In our case, the randomising and sub-setting were done 10,000 times. The observed absolute difference in means between the two groups is compared against this null distribution to arrive at a p-value. The p-value here is the probability of getting a difference in means as extreme as the observed absolute difference by chance alone.
